# Configurable Digital Virus Counter on Robust Universal DNA Chips

**DOI:** 10.1101/2020.10.22.350579

**Authors:** Elif Seymour, Nese Lortlar Ünlü, Eric P. Carter, John H. Connor, M. Selim Ünlü

**Affiliations:** Department of Biomedical Engineering, Boston University, Boston, Massachusetts 02215, United States; Department of Microbiology, Boston University School of Medicine, Boston, Massachusetts 02218, United States; Department of Electrical and Computer Engineering, Boston University, Boston, Massachusetts 02215, United States

**Keywords:** Imaging biosensor, DNA-directed antibody immobilization, homogeneous virus tagging, multiplexed detection, point-of-care diagnostics

## Abstract

Here, we demonstrate real-time multiplexed virus detection by applying DNA-directed antibody immobilization technique to a single-particle interferometric reflectance imaging sensor (SP-IRIS). In this technique, the biosensor chip surface spotted with different DNA sequences is converted to a multiplexed antibody array by flowing antibody-DNA conjugates and allowing specific DNA-DNA hybridization. The resulting antibody array is shown to detect three different recombinant Vesicular Stomatitis Viruses (rVSVs) genetically engineered to express surface glycoproteins of Ebola, Marburg, and Lassa viruses in real-time in a disposable microfluidic cartridge. We also show that this method can be modified to produce a single-step, homogeneous assay format by mixing the antibody-DNA conjugates with the virus sample in solution phase prior to flowing in the microfluidic cartridge, eliminating the antibody immobilization step. This homogenous approach achieved detection of the model Ebola virus, rVSV-EBOV, at a concentration of 100 PFU/ml in 1 hour. Finally, we demonstrate the feasibility of this homogeneous technique as a rapid test using a passive microfluidic cartridge. A concentration of 10^4^ PFU/ml was detectable under 10 minutes for the rVSV-Ebola virus. Utilizing DNA microarrays for antibody-based diagnostics is an alternative approach to antibody microarrays and offers advantages such as configurable sensor surface, long-term storage ability, and decreased antibody use. We believe these properties will make SP-IRIS a versatile and robust platform for point-of-care diagnostics applications.

---

Rapid and sensitive detection of viral infections is of significant importance for improving patient care and containing outbreaks that threaten public health. Current techniques employed in clinical diagnosis of viral infections include polymerase chain reaction (PCR), enzyme-linked immunosorbent assay (ELISA), isothermal nucleic acid amplification techniques (e.g. LAMP and RPA), or virus isolation in cell culture.^1,2^ These tests often require sending patient samples to a central laboratory with necessary equipment and trained personnel, and can take on the order of days, or weeks in the case of virus isolation from cell culture, hampering the fast containment of the virus and delaying the appropriate course of treatment. This situation is exacerbated when there is an excessive number of samples to be tested in an epidemic. Epidemics are one of the challenging problems that have caused widespread deaths since the beginning of the known human history. Starting from the post-classical era, humankind faced several epidemics such as the plague,^3^ viral hemorrhagic fever,^4^ cholera,^5^ smallpox,^6^ measles,^7^ poliomyelitis,^8^ and influenza.^9^ It is estimated that more than 20 million people died from Spanish flu (Swine flu H1N1) from 1918 to 1920.^10^ The rapid spread of the novel coronavirus, SARS-CoV-2, has reminded us that such pandemics are not historical anecdotes and exposed the major gaps in today’s infectious disease diagnostics. In many countries, due to the huge demand in RT-PCR tests, clinical laboratories have faced shortages of trained personnel, test reagents and lab space.^11^ The system has been rapidly overwhelmed, causing longer wait times and delaying appropriate isolation procedures. To ease the burden on centralized laboratories and ramp up the testing capacity, rapid point-of-care (POC) tests in lateral assay format have been developed. Although these tests can provide fast and simple detection, they lack sensitivity to detect low viral loads at the early stages of infection, when detection is essential for stopping community spread.^12^ There is a continuing need for alternative viral diagnostic techniques that can meet high sensitivity requirements in easy-to-use and portable POC platforms without the need for laboratory environment and trained personnel. An ideal POC platform should offer rapid (sample-to-answer in about 20 min) and sensitive detection with minimal sample preparation. Moreover, it should have multiplexing capability, which is especially important when it is necessary to distinguish between different viral pathogens that cause similar physical symptoms. It is also desirable that the diagnostic platform is easily configurable to new emerging viral pathogens with highly scalable and long-shelf-life consumables. In this paper, we describe a configurable and multiplexed digital virus counter capable of enumerating virions captured on a universal DNA sensor chip and its application as a rapid testing platform utilizing DNA-conjugated antibodies and disposable microfluidic cartridges.

Our group developed a label-free biosensor termed Interferometric Reflectance Imaging Sensor (IRIS) that has been used to quantify biomass accumulation on a microarray chip and was shown to detect virus particles captured onto an antibody-printed chip with a limit-of-detection (LOD) of 5×10^5^ PFU/ml.^13,14^ IRIS utilizes a silicon-silicon dioxide microarray chip and an imaging sensor, consisting of 4 different wavelength LEDs for illumination and a CCD camera that records reflected light intensities. Although, this sensor provided a simple, inexpensive, and high-throughput platform for virus detection, due to the ensemble-based nature of this biosensor, sensitivity of detection was moderate. The IRIS system was modified to image single virus particles and termed Single Particle – IRIS (SP-IRIS). SP-IRIS has been shown to individually count and size the nanoparticles bound to capture probes on the sensor surface over a large sensor area.^15^ In this technique, high affinity capture probes are immobilized on the surface that can selectively bind to the target virus. When virus particles bind to the surface, scattered light from the particles interfere with the reference beam reflecting from the layered substrate, allowing an enhanced signal from the particle that is detected on the CCD camera. Particles captured on the sensor surface appear as bright dots in the resulting image. This technique can also be used for Single-Particle Interferometric (SPIR) microscopy modality and provide shape and size information allowing detailed morphological characterization of viruses.^16^

SP-IRIS has been utilized for detecting viruses in complex media using antibody microarrays by imaging chips both dry (dried after sample incubation and wash steps) and in liquid using disposable microfluidic cartridges.^17–19^ Microfluidic integration of SP-IRIS provided an enclosed chamber for virus incubation, eliminated wash and drying steps and further improved the sensitivity of virus detection. These recent developments rendered SP-IRIS highly sensitive, fast and easy-to-use. (See Table S-1 of the Supporting Information for a comparison of different modalities of IRIS in terms of key sensor properties.) As we improved SP-IRIS to develop it as a robust POC diagnostic platform, we focused on optimizing virus capture efficiency of antibody microarray chips, one of the most important factors that affect assay sensitivity in solid-phase immunoassays. A major challenge with antibody-based solid-phase biosensors is the immobilization of capture probes on the sensor surface. The surface attachment chemistry can affect the biological activity of the antibody, its affinity and the background noise, ultimately affecting the sensitivity of the biosensor.^20–22^ Moreover, printing of antibodies on the microarray surface can introduce issues such as non-uniform surface coverage and assay-to-assay variability, affecting the assay accuracy and reproducibility.^23–25^

A DNA-based site-specific antibody immobilization technique, known as DNA-directed immobilization (DDI), has gained interest due to the reproducible production of DNA microarrays, compatibility of DNA microarrays with the fabrication of integrated microfluidic systems, and stability and robustness of DNA chips. In this technique, a universal ssDNA chip is first converted to an antibody microarray using antibodies tagged with short DNA sequences complementary to the immobilized DNA capture probes (Figure 1). The resulting DDI-antibody microarray can be used for detection of target using a variety of labeled or label-free biosensing techniques. DDI technique has been shown to improve the antigen binding capacity,^26,27^ the antibody surface coverage, and assay reproducibility,^28,29^ compared to directly immobilized antibodies. In a previous study, we applied DDI approach to SP-IRIS to demonstrate the label-free detection of whole viruses. We showed that DDI elevates the antibodies from the sensor surface and improves the virus capture efficiency, increasing the sensitivity of the SP-IRIS platform.^27,30^ Here, we extend this approach to show multiplexed detection of three virus pseudotypes genetically engineered to express Ebola, Marburg, and Lassa glycoproteins as a model for Ebola, Marburg and Lassa virus detection. Utilizing antibody - DNA conjugates to convert a DNA chip into a multiplexed antibody array suggests an alternative approach for generation of robust and repeatable diagnostic platforms by making use of stability and highly reproducible nature of DNA microarrays. We also demonstrate, for the first time, a homogenous DNA-directed virus capture assay where the antibody - DNA conjugates and the virus sample are mixed in solution phase before incubating the DNA chip. We present the combined utility of this homogeneous assay with a passive flow cartridge as an example of the application of SP-IRIS platform to a rapid test format which is suitable for POC testing.

**Figure 1:**
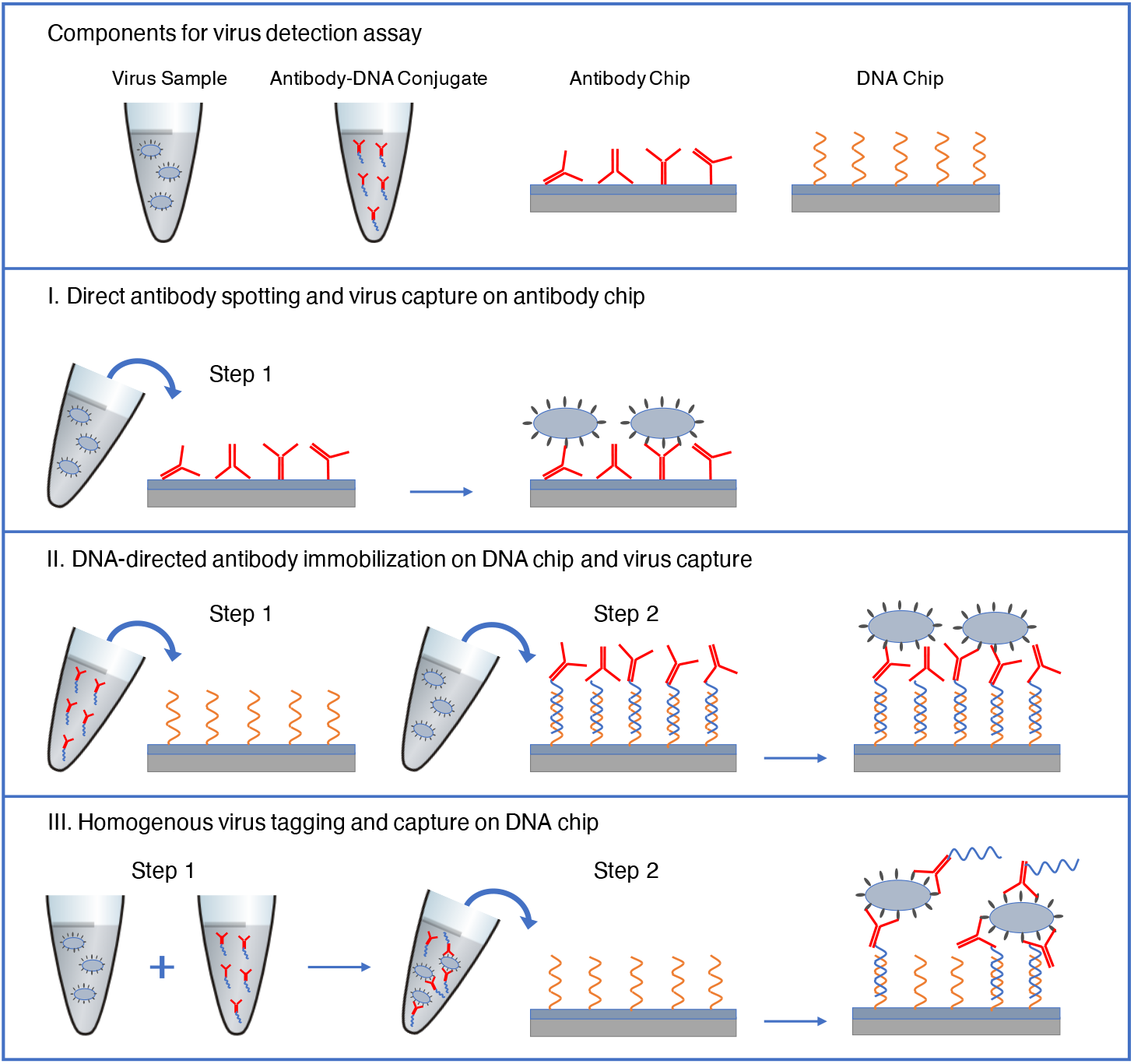
Three different approaches for virus capture on the SP-IRIS chip surface. I. Direct antibody spotting: Antibodies are immobilized on the polymer-coated chip surface directly and viruses are captured on the antibody surface. II. DNA-directed antibody immobilization: A ssDNA surface is converted to an antibody surface by an incubation step with antibody-DNA conjugates. III. Homogenous virus tagging in solution with antibody-DNA conjugates: Virus sample is first mixed with antibody-DNA conjugates allowing antibody-antigen binding in solution. This mixture is then applied to a ssDNA surface and DNA molecules protruding the viruses are captured on the surface through sequence specific DNA hybridization.

## METHODS

### Materials and Reagents

Silicon chips with a patterned thermally grown silicon dioxide were purchased from Silicon Valley Microelectronics Inc. An oxide thickness of 30 nm was used since the optimization studies for in-liquid visualization of the viruses showed that this thickness gave the highest level of particle contrast. Custom-designed, disposable, active and passive microfluidic cartridges were purchased from Aline. Monoclonal antibodies (mAbs) against Ebola virus glycoprotein (13F6),^31^ Marburg virus glycoprotein (AGP74-1),^32^ and Lassa virus glycoprotein (8.9F) were provided by Mapp Biopharmaceutical, Prof. Ayato Takada (Hokkaido University), and Prof. James Robinson (Tulane University), respectively. Recombinant Vesicular Stomatitis Virus (rVSV) stocks expressing surface glycoproteins of Ebola, Marburg, and Lassa viruses were created as described previously.^27^ HPLC purified 5’-aminated single-stranded DNA (ssDNA) molecules were purchased from Integrated DNA Technologies. Antibody-DNA conjugation kit was purchased from Innova Biosciences. Polymer kit for chip surface coating (MCP-2) was purchased from Lucidant Polymers.

### Sensor Chip Functionalization

The silicon-silicon dioxide chips were cleaned by sonicating in acetone and then rinsing with methanol and Nanopure water. Chips were then dried under nitrogen. Chips were coated with a 3-D polymeric coating, copoly(N,N-dimethylacrylamide (DMA) - acryloyloxysuccinimide (NAS) - 3(trimethoxysilyl) - propylmethacrylate (MAPS)) polymer, that offers a simple, inexpensive, and repeatable coating process and provides high density probe immobilization due to its 3-D structure.^33,34^ Copoly(DMA-NAS-MAPS) has NHS esters for covalent binding of proteins and amine-tagged DNA molecules. For coating, the chips were first treated with oxygen plasma and then immersed in 1× MCP-2 polymer solution for 30 min. Chips were then rinsed extensively with Nanopure water and dried with nitrogen. Polymer coated chips were baked at 80°C for 15 min and stored in a desiccator until microarray printing.

### Printing of Biomolecules on Sensor Chips

Antibody and DNA molecules were printed on the polymer-coated chips using a piezo-driven, non-contact dispensing system, sciFLEXARRAYER S3 (Scienion, Germany). 5’-aminated ssDNA surface probes were spotted at 30 μM in 150 mM sodium phosphate buffer (pH = 8.5), producing DNA spots of ~100 μm diameter. For passive cartridge experiment and stability test, 3 mg/ml 13F6 antibody in PBS with 50 mM Trehalose was spotted on the chip along with ssDNA (A’ sequence in Table 1). Antibody spots were ~150 μm in diameter. During spotting, humidity was kept at 58% in the spotter chamber and the spotted chips were kept in the chamber overnight at 67% humidity. The chips were then washed with 50 mM Ethanolamine in 1× Tris - buffered saline (150 mM NaCl and 50 mM Tris - HCl, Fisher Scientific), pH = 8.5, for 30 min to quench the unreacted NHS groups in the polymer. This step was followed by a 30 min wash with PBST (PBS with 0.1% Tween) and a rinse with PBS and Nanopure water. The chips were finally dried with nitrogen.

**Table 1:**
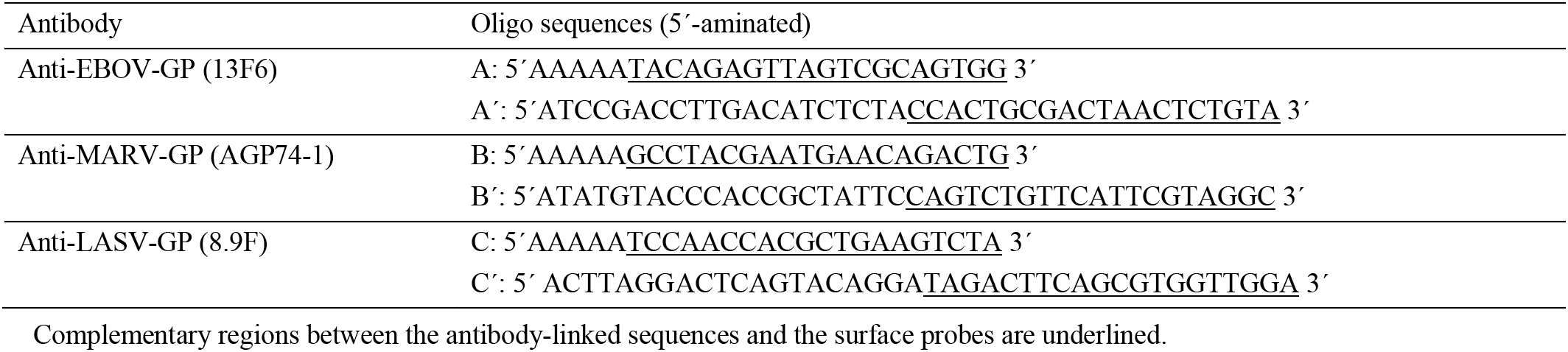
DNA sequences conjugated to the mAbs (A, B, and C) and partially complementary surface probes (A’, B’, and C’)

### Antibody-DNA Conjugation

Antibody - DNA conjugates were prepared using Thunder - Link Oligo Conjugation Kit (Innova Biosciences). Each monoclonal antibody (13F6, AGP74-1, and 8.9F, 1 mg/ml) was reacted with a specific 5’-aminated 25mer ssDNA (40 μM) according to the manufacturer’s instructions. The DNA concentration used in the conjugation was optimized to yield a DNA to antibody ratio between 1-2 in the final conjugate. 5’-aminated 40mer DNA sequences that are immobilized on the sensor chips are partially complementary to the antibody-conjugated DNA strands. Length of the DNA surface probes were optimized in a previous study that showed 40mer probes provide optimal elevation of the antibodies from the sensor surface. Three antibody-linked DNA sequences (A, B, and C) and corresponding surface-immobilized probe sequences (A’, B’, and C’) are given in Table 1. DNA sequences were designed by using OligoAnalyzer tool (Integrated DNA Technologies) to eliminate the formation of hairpin and self-dimer structures and to prevent cross hybridization between DNA sequences. A 5’ spacer sequence (5-bp polyA) was added to the antibody-linked DNA sequences to increase the hybridization efficiency.

As given by the Bradford assay for the protein part of the conjugate and the absorbance at 260 nm for the DNA part, DNA-to-Ab ratios were measured as 1.5, 1.7, and 1.5, for 13F6, AGP74-1, and 8.9F mAb conjugates, respectively. Antibody - DNA conjugates are designated in the text by adding the letter representing the DNA sequence to the antibody name as follows: anti-EBOV-DNA ‘A’, anti-MARV-DNA ‘B’, and anti-LASV-DNA ‘C’.

### Optical Biosensor Setup and Data Analysis

In-liquid virus detection experiments were performed using SP-IRIS and spotted biosensor chips mounted into either a multi-layer laminate, disposable, active flow cartridge or a disposable passive flow cartridge that is composed of laminate layers and an absorbent pad (Figure 2). For the active flow cell, the flow was controlled with a syringe pump (Harvard Apparatus, PHD 2000) and a flow rate of 3 μl/min was used.

**Figure 2:**
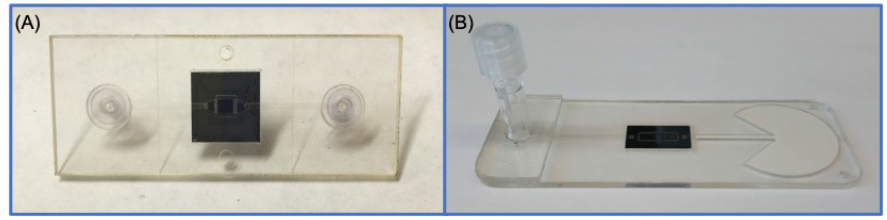
Active and passive flow microfluidic cartridges used for in-liquid virus detection experiments. (A) Active flow cartridge with fluid inlet and outlet designed to flow the sample over the SP-IRIS chip (10 mm x 10 mm). The channel is 25 μm-thick and 2 mm-wide. (B) Passive, adsorbent-pad integrated cartridge for capillary flow of the sample placed in the fluid reservoir.

SP-IRIS setup is composed of a single wavelength LED (525 nm) for illumination of the substrate, a high-magnification objective (40×, 0.9 NA) to obtain a high spatial resolution image, and a CCD camera. An autofocus system (MFC-2000, Applied Scientific Instrumentation) was used to control the focus during image acquisition. The recorded SP-IRIS images are analyzed for the bound virus particles for each spot. Image analysis is performed using a custom software that identifies the particle-associated intensity peaks in a given image and applies a Gaussian filter to eliminate the noise from the background. The morphological features of the antibody spots that become prominent due to the high resolution of the optical system cause a low correlation with a Gaussian-type intensity profile, and therefore, the background signal caused by these features is eliminated by adjusting the Gaussian filter parameters. SP-IRIS uses a forward-model to correlate the background normalized intensities of the particles to the particle size,^15^ allowing size-based filtering of the images to increase the specificity of detection. To quantify the virus particles in a spot, the diffraction-limited particles in the appropriate size range are detected and counted. The signal is expressed as virus density (number of particles per mm^2^) for a given spot by dividing the number of the detected particles by the analyzed spot area. For the end-point experiments, the initial particle count is subtracted from the final particle count for each spot to obtain the net number of particles bound to the spot during the experiment.

### Real-time Multiplexed Virus Detection in Active Microfluidic Cartridge

Three ssDNA surface probes (A’, B’, and C’ in Table 1) that are partially complementary to the antibody-linked DNA sequences were spotted on a polymer coated SP-IRIS chip at a 30 μM concentration. The chip was mounted in the active microfluidic cartridge via a pressure sensitive adhesive (PSA) and the assembled cartridge was placed on the SP-IRIS stage. First, a mixture of DNA-conjugated anti-EBOV, anti-MARV, and anti-LASV mAbs (at 5 μg/ml in PBS with 1% BSA) was flowed through the channel for 30 min at a rate of 3 μl/min. After a 400 μl wash step with PBS, recombinant VSV models of EBOV, MARV and LASV were flowed sequentially over the SP-IRIS chip, by flowing each VSV pseudotype for 30 min. 400 μl PBS was flowed through the channel after each virus incubation to wash the extra virus in the channel and the tubing. The order of the virus incubation was rVSV-EBOV, rVSV-MARV, and rVSV-LASV, and their titers were 10^4^, 10^4^ and 10^5^ PFU/ml, respectively, as determined by the plaque assay. The images of the anti-EBOV, anti-MARV, and anti-LASV spots generated by DDI were acquired every minute, and the virus densities on each spot were calculated over the course of the experiment.

### LOD Determination for One-step Homogeneous Detection of rVSV-EBOV

One-step homogeneous assay uses a DNA chip and a solution-phase mixture of the virus sample and antibody-DNA conjugates, eliminating the antibody-DNA conjugate incubation step. To determine the LOD for the homogenous, DNA-directed rVSV-EBOV assay using SP-IRIS, we performed a dilution experiment with 10-fold dilutions of a 10^6^ PFU/ml rVSV-EBOV stock, ranging from 10^5^ PFU/ml to 10^2^ PFU/ml. Five SP-IRIS chips were spotted with 6 replicates of A’ probe and washed as described previously. 1 μl of anti-EBOV-DNA ‘A’ conjugate at 50 μg/ml was mixed with 0.5 ml of each of the rVSV-EBOV dilutions prepared in 0.1× PBS with 1% BSA. A blank sample was also prepared by mixing the same amount of Ab-DNA conjugate with 0.5 ml 0.1× PBS with 1% BSA. After waiting for 15 min, 180 μl of each virus dilution and the blank sample was passed over a different SP-IRIS chip in the active microfluidic cartridge in subsequent experiments. For each sample, the channel was first filled with 0.1× PBS with 1% BSA and the spots were scanned to obtain the pre-incubation particle counts. Then, the sample was flowed for 1 h in the cartridge at a rate of 3 μl/min. After the channel was washed with PBS, the spots were scanned again. The net number of virus particles captured on the A’ spots were counted and the average virus densities were calculated from 6 replicate spots for each chip.

### Combining Homogenous DNA-directed Assay with Passive Microfluidic Cartridge

Passive microfluidic cartridge has been designed to simplify the SP-IRIS platform by eliminating the need for an active syringe pump and to create a fully-contained test platform in order to minimize the sample handling.^19^ Briefly, the passive microfluidic cartridge consists of a sample reservoir with a vented luer cap and an integrated 270°C fan shape absorbent pad in the channel placed after the chip (Figure 2). The sample to be tested is pipetted into the reservoir and the flow is established by applying a pressure through the closure of the reservoir cap. Once the sample flows over the chip and touches to the absorbent pad on the other side, the adhesive sealing tab on the cap is removed to let the fluid migrate under the atmospheric pressure. A stable flow rate (~3 μl/min) is established by the 270°C fan shape of the absorbent pad.

To demonstrate the feasibility of using the homogeneous assay in combination with the passive flow cartridge and to compare its performance to the directly immobilized antibody assay, an SP-IRIS chip was printed with anti-EBOV mAb, A’ probe, and a negative DNA sequence, and washed as described previously. 0.5 μl of 50 μg/ml anti-EBOV-DNA ‘A’ conjugate was added to 250 μl of 10^4^ PFU/ml rVSV-EBOV sample in PBS with 1% BSA. After 5 min incubation, 100 μl of this mixture was placed in the sample reservoir of the passive microfluidic cartridge. The reservoir cap was closed and tightened until the liquid started touching the absorbent pad. The cartridge was immediately placed on the SP-IRIS stage and the images of the directly immobilized anti-EBOV, A’ probe, and negative DNA spots were recorded every 2 min during a 30 min incubation. Following the image acquisition, the virus density was calculated for each of the three spot types at every time point to show the real-time binding of the viruses.

### Accelerated Stability Testing of Dried Antibody-DNA Conjugates

3 μl of 50 μg/ml anti-EBOV-DNA ‘A’ was aliquoted into five tubes and placed in a vacuum oven for 30 min at 37°C for drying the conjugate solution. After drying, 4 of the dried conjugate tubes were placed into the oven at 35°C for the accelerated stability test. One tube was used for the Day 0 measurement on the same day. SP-IRIS chips, spotted with 6 replicates of anti-EBOV antibody and A’ sequence, were also kept in the oven at 35°C. 10^4^ PFU/ml rVSV-EBOV sample was prepared on Day 0 and aliquoted into five tubes. Four of the virus samples were stored at −80°C until the virus detection experiments. On each of the days 0, 3, 7, 10, and 14, one dried conjugate tube was reconstituted with 100 μl PBS with 1% BSA and flowed over the SP-IRIS chip mounted on the active microfluidic cartridge for 30 min for DDI. The channel was washed with PBS and the spots were imaged with SP-IRIS for the pre-incubation particle counts. Then, 10^4^ PFU/ml rVSV-EBOV sample was flowed over the chip for 30 min and the spots were scanned again to obtain the post-incubation particle counts. Average virus densities on the DDI-antibody spots were calculated from 6 replicate spots for each chip.

## RESULTS & DISCUSSION

### Multiplexed, Real-time Detection of rVSV-EBOV, rVSV-MARV, and rVSV-LASV Using DDI-based SP-IRIS

To show the multiplexed detection of the rVSV models for Ebola, Marburg, and Lassa viruses and specificity of the antibody-DNA conjugates, we performed a sequential incubation with these viruses after functionalizing a DNA spotted SP-IRIS chip with three antibody-DNA conjugates. First, a mixture of DNA conjugated anti-EBOV, anti-MARV, and anti-LASV mAbs was flowed through the active microfluidic cartridge for 30 minutes. This step loaded the antibodies onto the specific complementary ssDNA spots on the sensor chip. Following the antibody immobilization step, rVSV samples were flowed sequentially in the following order: rVSV-EBOV, rVSV-MARV, and rVSV-LASV. Each virus sample was flowed for 30 minutes followed by a 400 μl PBS wash step. SP-IRIS image acquisition was done with 1 min intervals. Figure 3 shows the virus densities (particle count / mm^2^) on the three DDI-antibody spots (anti-EBOV-DNA ‘A’, anti-MARV-DNA ‘B’, and anti-LASV-DNA ‘C’) during the course of the experiment. Following the rVSV-EBOV sample addition (red band), the signal on the anti-EBOV-DNA ‘A’ spot starts to increase whereas the other two spots do not show any virus binding. After the rVSV-MARV (green band) is introduced, the virus density on the anti-MARV-DNA ‘B’ spot starts to increase showing the specific detection of rVSV-MARV. Finally, when the rVSV-LASV sample is flowed in the channel (blue band), the virus density on the anti-LASV-DNA ‘C’ spot increases whereas the signal on the other two spots remain constant. Overall, these results show that the site-specific self-assembly of the three antibody – DNA conjugates on a DNA surface was performed successfully to generate a multiplexed antibody microarray, and each antibody-DNA conjugate was able to detect its target virus specifically with no cross-reactivity from other viruses. Such a programmable DNA surface can serve as a universal chip that can be adapted for the detection of different target viruses by using different sets of antibody-DNA conjugates based on the need.

**Figure 3:**
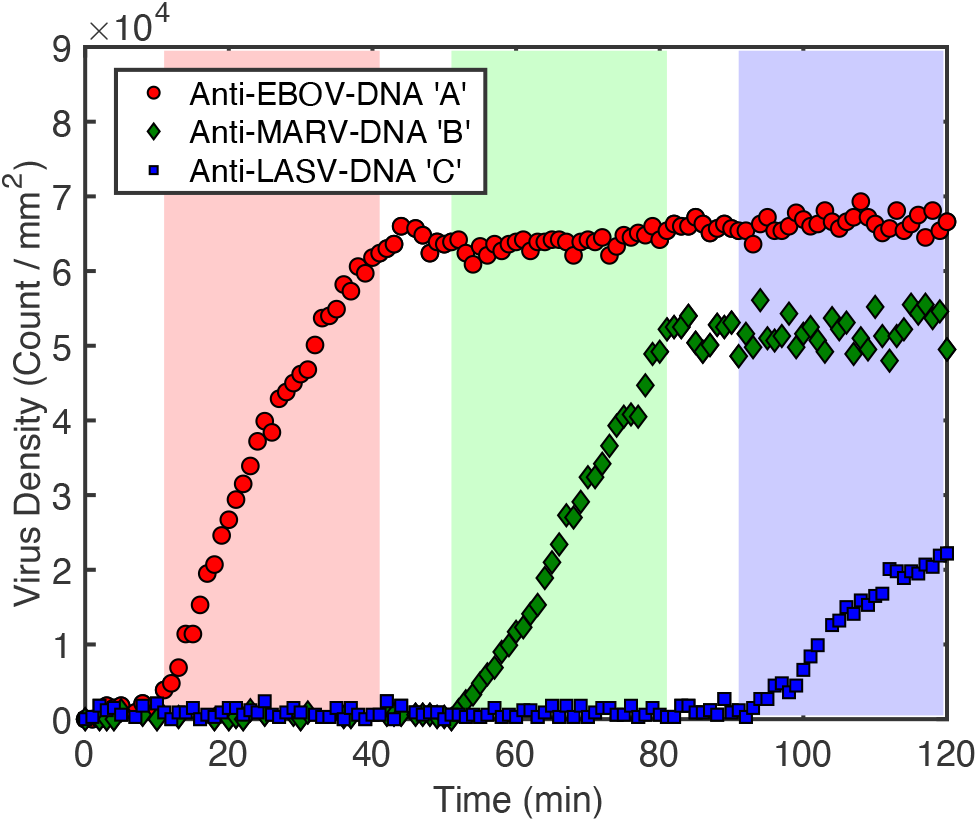
Multiplexed detection of the three different rVSVs for Ebola, Marburg and Lassa virus models on SP-IRIS using DNA-directed self-assembly of antibody-DNA conjugates. First, the SP-IRIS chip was incubated with a mixture of three different antibody-DNA conjugates that are specific for the surface glycoproteins of Ebola, Marburg, and Lassa viruses. Following the multiplexed antibody surface generation, rVSV-EBOV (red band), rVSV-MARV (green band), and rVSV-LASV (blue band) samples were flowed over the chip sequentially, with each incubation being 30 minutes. Virus densities for three DDI-antibody spots are shown in real-time, with 1 min intervals.

### Homogeneous Tagging of Ebola Virus Model with Antibody-DNA Conjugates in Solution-phase

One drawback of the conventional DDI-based detection assay is the time associated with the antibody-DNA conjugate immobilization step. To reduce the assay time and make this approach compatible with passive flow platforms where it is not practical to have sequential flows, we explored a homogeneous tagging approach where the virus sample is mixed with the antibody-DNA conjugates in solution prior to the incubation of the chip. This approach replaces the 30-min antibody immobilization step of the conventional DDI technique with a simple and fast mixing step, reducing the number of incubation and wash steps. Although the homogenous tagging of the target has been demonstrated for the detection of antigens previously,^28,35^ our work is the first one, to the best of our knowledge, to show the capture of whole viruses decorated with DNA-encoded antibodies on a ssDNA microarray surface.

One important consideration that needs to be addressed for this approach is the amount of the antibody-DNA conjugates to be added to the virus sample. Presence of excess antibody-DNA conjugates in the solution would cause blocking of the DNA surface with antibody-DNA conjugates, preventing the binding of the virus particles that are already decorated with antibody-DNA conjugates. We found that a concentration of 0.1 μg/ml antibody-DNA conjugate does not saturate the surface, causing only 0.5 nm height increase, as measured by IRIS (data not shown), compared to about a 4 nm height increase when the surface is fully saturated with the antibody-DNA conjugates. (In IRIS, 1 nm surface height corresponds to a surface antibody density of 1.2 ng/mm^2^.) We mixed the anti-EBOV-DNA ‘A’ conjugate (at a final concentration of 0.1 μg/ml) with the rVSV-EBOV samples prepared by 10-fold dilutions from a 10^6^ PFU/ml stock, ranging between 10^5^ - 10^2^ PFU/ml, and waited for 15 min prior to the flow over the SP-IRIS chip. Virus titer of the rVSV-EBOV stock was measured by plaque assay. Next, we flowed this mixture over the SP-IRIS chip in the active microfluidic cartridge for 1 h and determined the captured virus density on the complementary A’ spots. Figure 4 shows the average virus densities on the A’ spots obtained from 6 replicate spots for each titer tested. The detection threshold, indicated by solid red line, was calculated as the average virus density of six A’ spots plus three times the standard deviation from the blank chip that was incubated with only anti-EBOV-DNA ‘A’ conjugate. Average virus density from the blank chip, which is 636 virus count/mm^2^, is also shown in the graph as dashed red line. In our earlier work, we have demonstrated that variance scales inversely with the sensor area (equivalently number of spots averaged).^36^ Thus, for a large sensor area (or many spots), the LOD will depend only on the mean virus density of the blank chip. Therefore, the LOD for a single spot detection is 1672 virus count/mm^2^ (solid red line) corresponding to 43 PFU/ml, obtained by the extrapolation of the linear fit based on the data presented in Figure 4. Similarly, for a large area sensor where many spots can be averaged to virtually eliminate the variance, the LOD is expected to approach 636 virus count/mm^2^, further improving the sensitivity. The LOD for the homogeneous assay (43 PFU/ml) is comparable to the one obtained from the heterogenous DDI technique, 53 PFU/ml,^27^ and therefore, our results indicate that the homogenous approach provides a simpler assay procedure without affecting the sensitivity of the detection.

**Figure 4:**
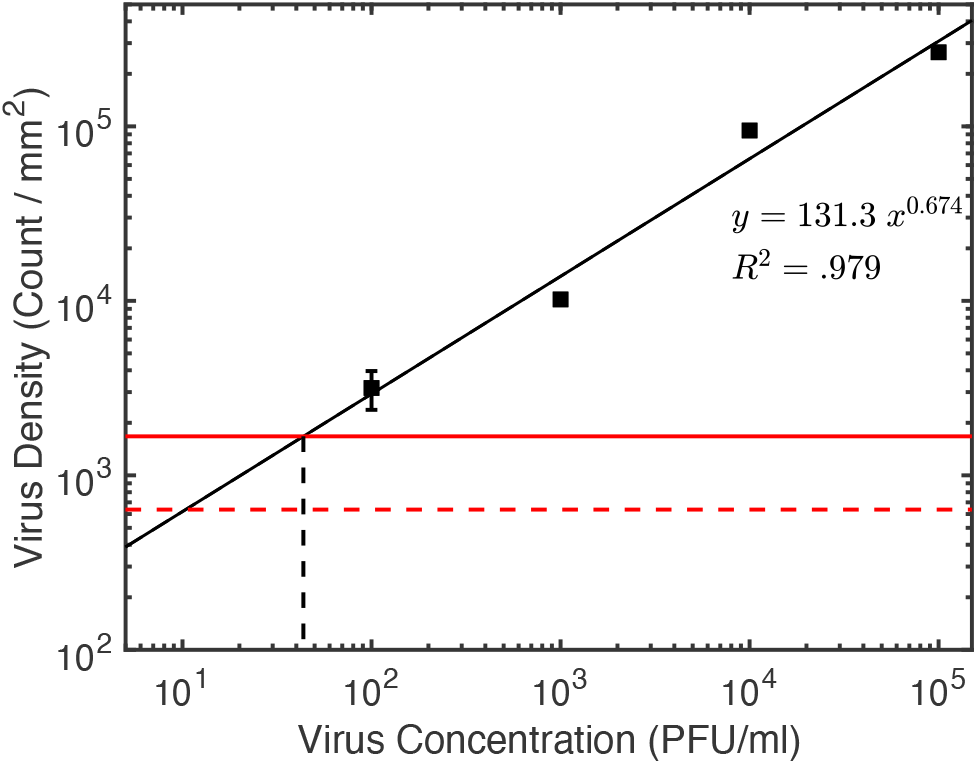
Dilution experiment using single - step homogeneous assay approach. Average virus densities are calculated from 6 replicate A’ spots that are complementary to the anti-EBOV-DNA ‘A’ conjugate. Solid red line shows the detection threshold calculated as the mean virus density from six spots plus three standard deviation of the mean from a blank chip. 100 PFU/ml rVSV-EBOV was detectable for a 1-hour incubation. LOD, based on the extrapolation of the linear fit, is 43 PFU/ml.

SP-IRIS platform provides a positive result once the adequate number of virions are counted, and therefore, test time can be significantly reduced for high titer samples. According to Figure 4, the experimental virion count response scales with [virus concentration] ^(2/3)^ - a perfect theoretical fit to the sheet density of the virus on the chip surface. In contrast, in an RT-PCR test, the cycle threshold (Ct) values are inversely and logarithmically related to the viral RNA copy number. Ct values reported for SARS-CoV-2 in a recent study are 30.76, 27.67, 24.56, and 21.48, corresponding to 1.5×10^4^, 1.5×10^5^, 1.5×10^6^, and 1.5×10^7^ copies/ml, respectively.^37^ While a thousand-fold increase in viral load provides a marginal (30%) reduction in Ct values corresponding to a small time saving for RT-PCR (about 10 min), the reduction in test time for SP-IRIS can be as high as 100-fold. Based on our data presented in this and next section, SP-IRIS is expected to have significantly reduced test times at median viral loads (a few minutes) allowing for high throughput testing.

One other advantage of the homogenous assay is the decreased antibody usage compared to the direct immobilization and the conventional DDI. Homogenous assay uses at least three-fold less antibody per test, decreasing the cost of the assay. (See Supporting Information for the comparison of the antibody usage for three methods.) Our LOD determination experiment used a high number of antibody-DNA conjugates per virus particle. This amount can be further decreased (at least 3-fold) and still have sufficient Ab-DNA molecules in the solution for efficient tagging of the virus particles (Table S-2 of the Supporting Information). Moreover, the use of low antibody concentrations makes testing of the antibodies possible even when they are available at low quantities or concentrations.

### Passive Microfluidic Cartridge Test Using the Homogeneous Approach

Active flow cartridge-based approach is not practical for field testing due to the complexity of the sample flow process that uses a syringe pump and tubing. This especially brings concerns in the case of lethal virus outbreaks due to the risk associated with the sample fluid handling. Therefore, we have designed a passive microfluidic cartridge that would eliminate the need for an active pump and provide a fully contained test platform with minimum sample handling.^19^ We combined the homogeneous virus tagging approach with this lateral-flow cartridge to show the utility of SP-IRIS as a rapid testing platform. For this purpose, we added the anti-EBOV-DNA ‘A’ conjugate (final concentration of 0.1 μg/ml) to the rVSV-EBOV sample (10^4^ PFU/ml), mixed, and waited for 5 minutes. Then, we applied this mixture to the reservoir and closed the cap. Once the flow started, we put the cartridge onto the SP-IRIS stage and started scanning the DNA spots (both complementary and negative DNA sequences) and directly immobilized anti-EBOV spots with 2 min intervals.

Figure 5 shows the captured virus density as a function of time for complementary DNA spots (A’), negative DNA spots and directly immobilized anti-EBOV antibody. A positive signal can be observed on the complementary DNA spots within the first 6 minute of the incubation. Moreover, the signal from the directly immobilized anti-EBOV spots is lower than that of DNA spots which is consistent with our previous findings. Our results suggest that the combination of the homogenous assay with the lateral-flow cartridge can provide a suitable platform for rapid and sensitive virus diagnostics.

**Figure 5:**
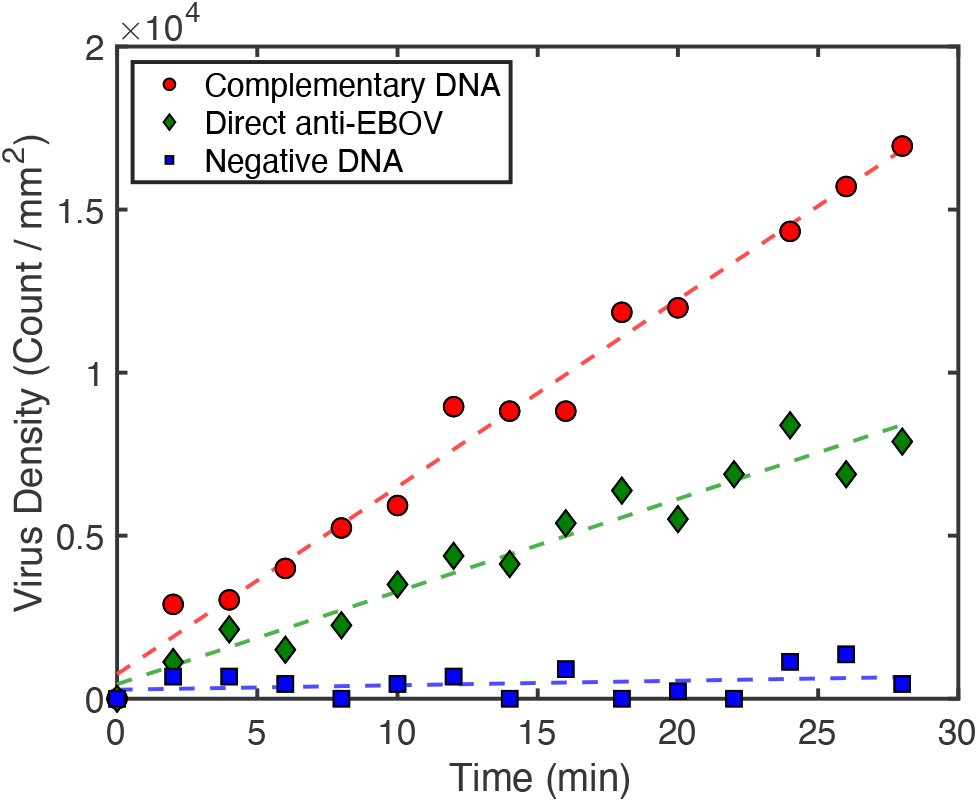
Real-time virus capture in a lateral-flow cartridge using one-step, homogeneous virus tagging approach. 10^4^ PFU/ml rVSV-EBOV was mixed with anti-EBOV-DNA ‘A’ conjugate, and this mixture was applied to the passive cartridge. Once the flow started, images were taken with SP-IRIS every 2 min for complementary DNA (A’), negative DNA, and directly immobilized antibody spots. A positive signal was detected as early as 6 minutes on the complementary DNA spots. Negative DNA spots, that are not complementary to the antibody-DNA conjugate, do not show any virus binding.

### Accelerated Stability Test with Dried Antibody-DNA Conjugate

To test the stability of the dried antibody-DNA conjugates, we performed an accelerated stability test by storing the dried anti-EBOV-DNA ‘A’ conjugates and the spotted SP-IRIS chips at 35°C for 2 weeks and performing in-liquid virus detection experiments using DDI on the days 0, 3, 7, 11 and 14. Figure 6 shows how the SP-IRIS signal changes over time for the DNA-conjugated antibodies that were reconstituted for DDI on the test day. At the end of the two weeks, the antibody-DNA conjugates captured approximately 2.5 times more viruses per mm^2^ than the directly immobilized antibody spots on the day 0 chip (shown by the dashed red line), showing the superior performance of DDI technique to the direct spotting. The signal on the directly spotted antibodies showed a faster degradation rate over time and higher variability compared to the dried antibody-DNA conjugates (data not shown). By extrapolating the line fit for the data points in Figure 6, the stability of the antibody-DNA conjugates is calculated as 59 days at 35°C. Using the Q-Rule ^38^, the shelf-life of dried antibody conjugates at 25°C is estimated to be 177 days (See Supporting Information for the details of the shelf-life calculation). Our results indicate that the dried Ab-DNA conjugates stay functional for extended periods of time without the need for refrigeration. The shelf-life of the antibody-DNA conjugates can be further improved by freeze-drying and using optimized excipient formulations.^39^

**Figure 6:**
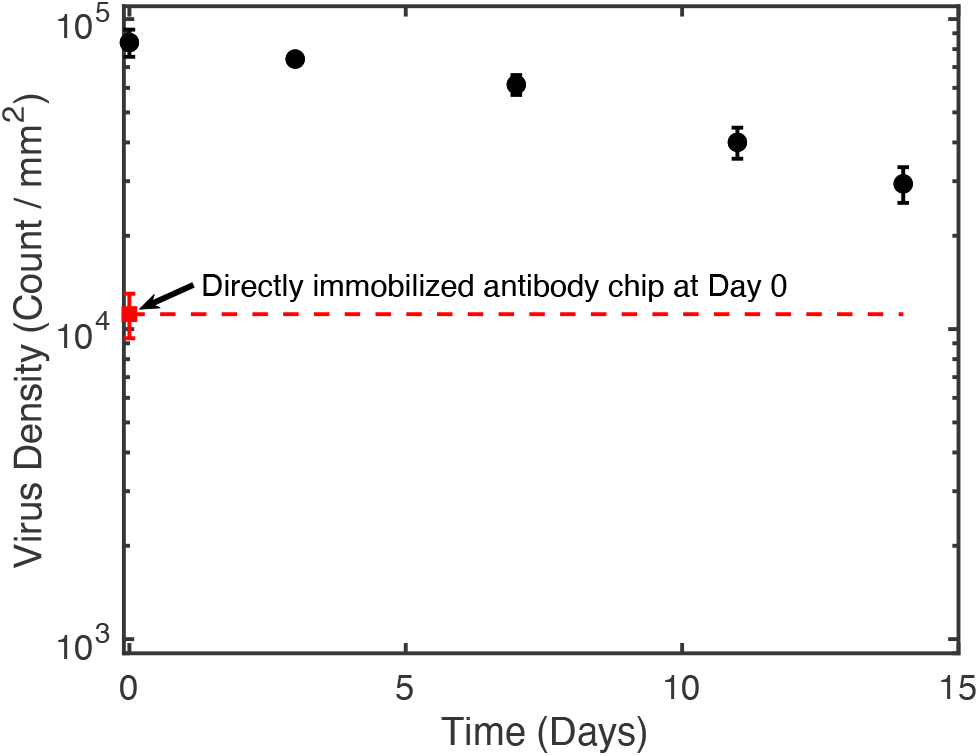
The accelerated stability testing of the dried antibody-DNA conjugates at 35°C over a 2-week period. Average virus densities on the DDI-antibody spots (n=6 replicate spots) are shown for the rVSV-EBOV detection experiments performed on the days 0, 3, 7, 10, and 14. Dashed red line represents the performance of the directly spotted anti-EBOV antibodies on the fresh chip at day 0.

## CONCLUSIONS

We applied the DDI technique to our virus counter, SP-IRIS, for generation of a multiplexed antibody array for the detection of Ebola, Marburg, and Lassa viruses. We showed the specific self-assembly of the antibodies on a DNA microarray surface and subsequent real-time detection of three different rVSVs in a disposable microfluidic cartridge. We also demonstrated the homogeneous tagging of the viruses with antibody-DNA conjugates in solution phase. By introducing this DNA-encoded virus tagging approach, antibody immobilization step in conventional DDI can be eliminated, decreasing the assay time and complexity. In addition, homogenous DNA-directed assay uses substantially less antibody (three-fold) compared to the conventional DDI and direct spotting. Moreover, in-solution binding of antibodies eliminates the problems that affect capture efficiency such as antibody orientation, steric hindrance, and the activity loss due to antibody immobilization. We also demonstrated the combined utility of this homogenous method with a passive microfluidic cartridge. This platform allowed us to detect 10^4^ PFU/ml rVSV-EBOV in less than 10 min in a disposable, contained cartridge, showing the feasibility of SP-IRIS as a rapid and sensitive viral diagnostic platform.

DNA chips also offer the advantage of being configurable, allowing the use of the same multiplexed DNA chips to create the desired virus detection panel. For example, clinicians would greatly benefit from a multiplexed test for SARS-CoV-2 and different influenza viruses to differentiate between these viruses that cause similar symptoms and that can happen concurrently. We envision that a multiplexed, DNA microarray-based SP-IRIS system would provide a versatile, robust, and simple diagnostic platform through the use of universal DNA chips and specific antibody-DNA conjugates that can be synthesized quickly according to the need.

## ASSOCIATED CONTENT

### Supporting Information

Comparing antibody usage for direct immobilization and DNA-directed assays, Table S-1: Comparison of IRIS platforms in terms of key POC biosensor properties, Table S-2: Number of antibody-DNA conjugates per virus particle used in a homogenous DNA-directed virus capture assay, Antibody-DNA conjugate shelf-life calculation

## ACKNOWLEDGEMENTS

Authors would like to thank Steven M. Scherr for his help with microfluidic cartridge experiments and A. J. Devaux for taking cartridge pictures. This work was supported by R01AI1096159 to J.H.C. and M.S.U.

