## Supporting Information for "Configurable Digital Virus Counter on Robust Universal DNA Chips"

**Comparing antibody use for direct immobilization and DNA-directed assays**

In one spotting run, 10 µl of a 3 mg/ml antibody is used. Nozzle that dispenses the liquid to the chips is allowed to withdraw 4-5 µl of this liquid to avoid bubble formation in the nozzle. One drop volume is approximately 0.2 nl, and two to three drops are used for each spot. Assuming that 2 drops are used for each spot, 12,500 spots can be generated from a 5 µl antibody solution. One 10 mm x 10 mm SP-IRIS chip, with an active spotting area of 2.3 mm x 2.3 mm, can have 64 spots on it. Therefore, 12,500 spot capacity would allow for preparation of 195 chips with 30 µg of antibody.

For conventional, two-step DDI-based assay, we used 500 ng antibody (100 µl of 5µg/ml Ab-DNA conjugate) to ensure surface saturation for optimized virus capture in the multiplexed virus detection experiment. However, as shown by the accelerated stability experiment results, 150 ng of the conjugate was sufficient to generate an antibody array with the same performance. Therefore, it is possible to run 200 tests with 30 µg of antibody by using 150 ng of the conjugate per test.

In homogeneous DNA-directed assay, one test (one chip) uses 50 ng antibody (1 µl of 50 µg/ml Ab-DNA conjugate), allowing 600 chips to be tested with 30 µg of antibody. Therefore, three times as many tests can be performed with the same amount of antibody when compared to the direct immobilization and the DDI techniques.

**Table S-1: Comparison of IRIS platforms in terms of key POC biosensor properties**

| | IRIS | SP-IRIS | In-liquid SP-IRIS | DDI based in-liquid SP-IRIS | Homogenous DDI based in-liquid SP-IRIS |
|---|---|---|---|---|---|
| LOD for whole virus detection | $3.5 \times 10^5$ PFU/ml [1] | $5 \times 10^3$ PFU/ml [2] | 645 PFU/ml [3] | 53 PFU/ml [3] | 43 PFU/ml (This work) |
| Sensor chip surface | Antibody | Antibody | Antibody | DNA | DNA |
| Multiplexing | ✓ | ✓ | ✓ | ✓ | ✓ |
| Disposable cartridge | x | x | ✓ | ✓ | ✓ |
| No wash steps required | x | x | ✓ | x | ✓ |
| Compatibility with a lateral flow cartridge | x | x | ✓ | x | ✓ |
| Number of tests that can be done using 30 ug antibody* | 195 | 195 | 195 | 200 | 600 |
| Availability of freeze-drying option | x | x | x | ✓ | ✓ |

*The quality control process of the spotted chips is not taken into account when calculating these numbers. For direct antibody spotting, it is expected that a higher percentage of chips would fail the quality control process compared to the DNA chips due to the variability of protein immobilization.

**Table S-2: Number of antibody-DNA conjugates per virus particle used in a homogenous DNA-directed virus capture assay**

| | Number of virus particles in 0.5 ml of virus sample (a factor of 25 is used for PFU to virus particle number conversion) | Number of antibody-DNA conjugates per virus |
|---|---|---|
| $10^8$ PFU/ml | $1.25 \times 10^9$ | 158 |
| $10^7$ PFU/ml | $1.25 \times 10^8$ | 1,584 |
| $10^6$ PFU/ml | $1.25 \times 10^7$ | 15,840 |
| $10^5$ PFU/ml | $1.25 \times 10^6$ | 158,400 |
| $10^4$ PFU/ml | $1.25 \times 10^5$ | 1,584,000 |
| $10^3$ PFU/ml | $1.25 \times 10^4$ | 15,840,000 |
| $10^2$ PFU/ml | $1.25 \times 10^3$ | 150,840,000 |

In Table S-2, the number of antibody-DNA conjugates per virus particle is given for different concentrations of virus sample for the homogenous virus tagging. Even at the highest virus concentration ($10^8$ PFU/ml), there are 158 antibody-DNA conjugate molecules per virus, which is sufficient for efficient virus tagging. We believe that the amount of the Ab-DNA conjugate per test can be further decreased (at least 3-fold) without affecting the sensitivity. This would allow at least 9-fold less antibody usage compared to the conventional antibody immobilization.

### Antibody-DNA Conjugate Shelf-life Calculation

We calculated the stability of the dried Ab-DNA conjugate at 35°C by extrapolating the line fit in Figure 6 and assuming an LOD of 1000 virus count/mm$^2$, which gave ~59 days. According to the *Q* rule, the stability at a given lower temperature is calculated as:

Stability at $T_1 = Q^n$ x (Stability at $T_2$), where $T_1 < T_2$ and,

$n$ = Temperature change in °C / 10°C

For our accelerated stability test, $n = (35-25) / 10 = 1$. The value of $Q$ is typically set at 2, 3, or 4. Assuming $Q = 3$, the stability of Ab-DNA conjugate at 25°C is calculated as:

Stability at 25°C = 3 x 59 = 177 days

Stability of the directly immobilized Ab chips at 35°C was calculated from the accelerated stability testing data (data not shown) as ~28 days. Therefore, the stability of conventional Ab chips at 25°C is:

Stability at 25°C = 3 x 28 = 84 days.
